## Supplementary Information for "Super-resolution microscopy of mitochondrial mRNAs"

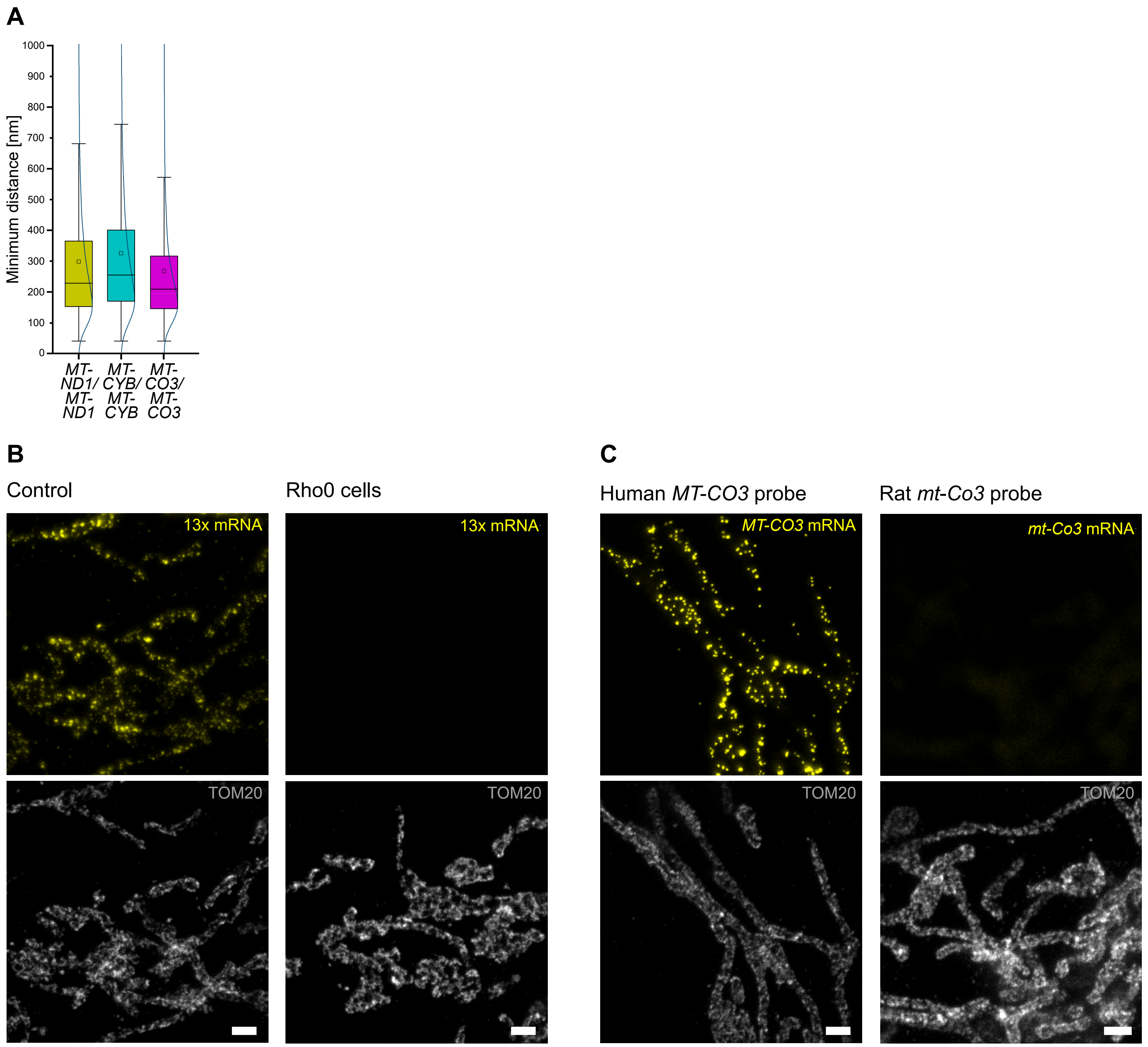


### Suppl Figure 1: STED-smFISH data self-distances and controls

**(A)** Quantification of the minimum self-distances of *MT-ND1*, *MT-CO3*, and *MT-CYB* mRNAs. Analysis is based on the same image data as Fig. 1F. **(B)** STED images of WT (Control, left) and Rho0 (right) U-2 OS cells labeled with probe pairs targeting all 13 mRNAs (yellow, top), together with an antibody targeting TOM20 (gray, bottom). No mRNA signal is present in mitochondria of Rho0 cells. **(C)** STED images of U-2 OS WT cells labeled with probe pairs targeting human *MT-CO3* mRNA (top left) or rat *mt-Co3* mRNA (top right) together with an antibody targeting TOM20 (gray, bottom). No rat *mt-Co3* mRNA signal is present in the human U-2 OS cells. Box: 25/75 percentile; whiskers: max/min without outliers; line: median; square: mean. Curved line: kernel density estimation representing the data point distribution (A). Scale bars: 1 µm (B, C).

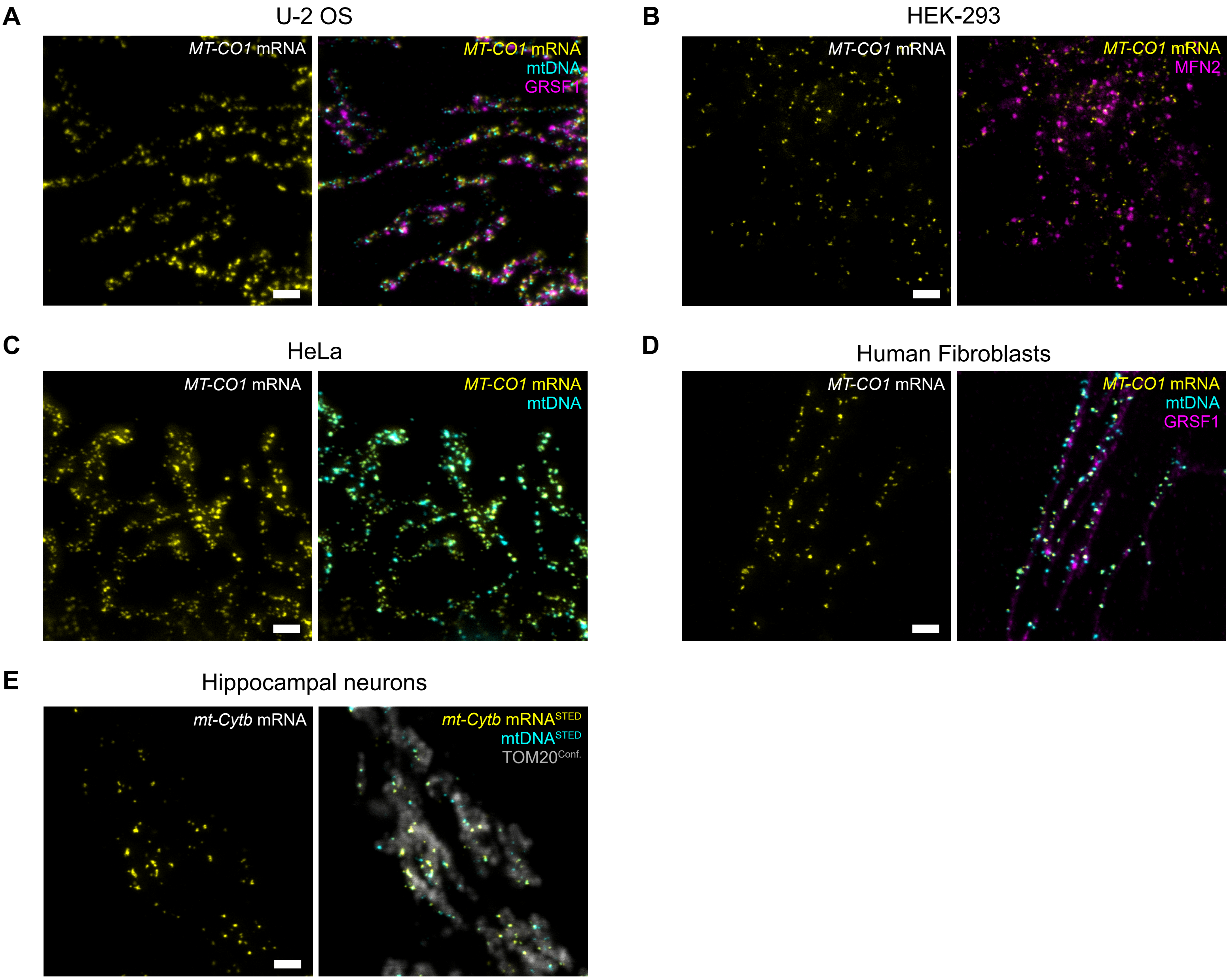


### Suppl Figure 2: STED-smFISH on different cell lines and primary neurons

The different cell types U-2 OS (A), HEK-293 (B), HeLa (C), primary human fibroblasts (D), and primary hippocampal neurons from rat (E) were labeled for *MT-CO1*-mRNA (A-D, yellow) and *mt-Cyb* mRNA (E, yellow), GRSF1 (A, D, magenta), mtDNA (A, C, D-E, cyan) and Mitofusin 2 (MFN2, B, magenta). Scale bars: 1 µm.

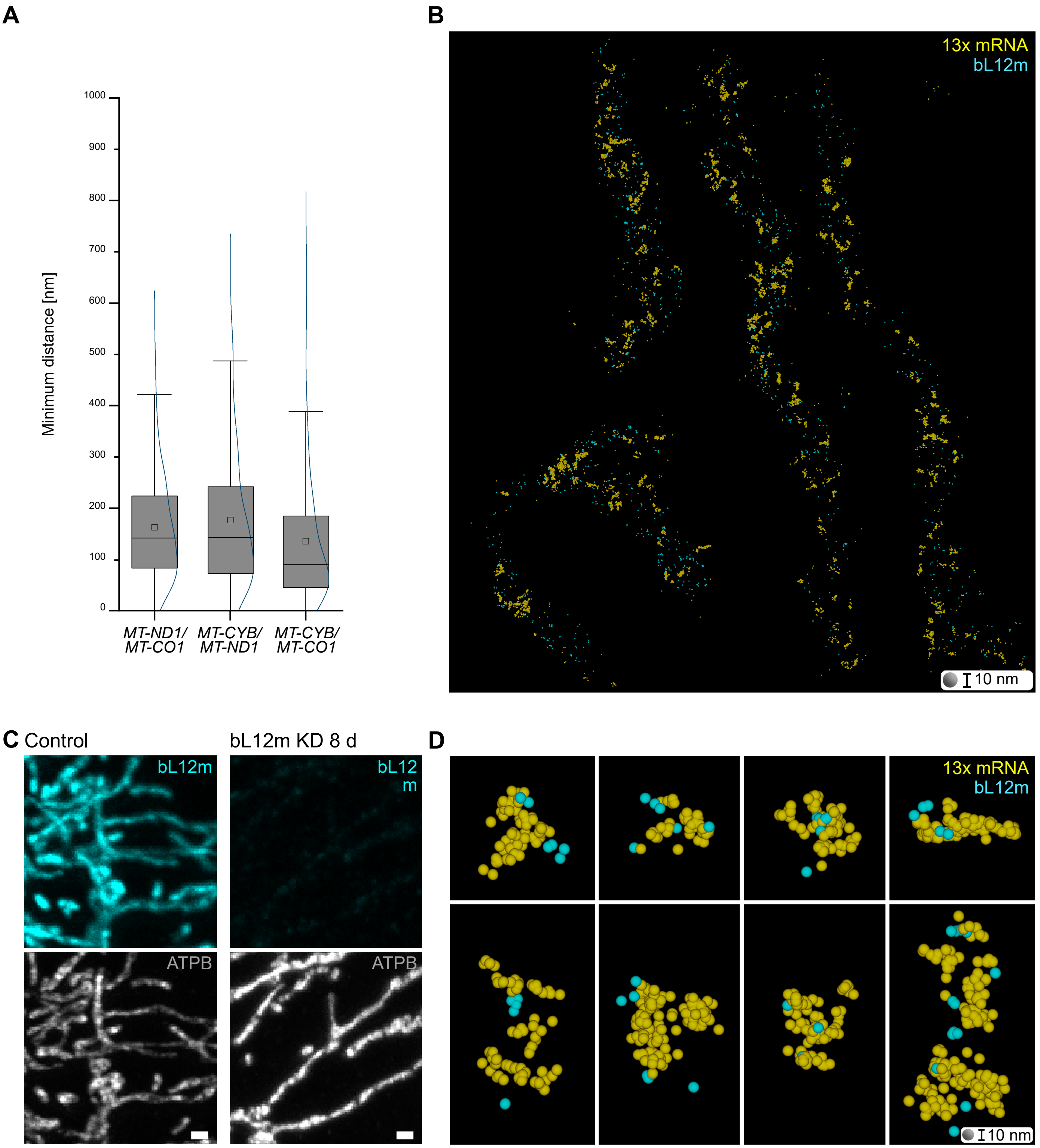


### Suppl Figure 3: MINFLUX-smFISH of all mRNAs and a ribosomal subunit

**(A)** Quantification of the minimum pairwise distances of emulated STED data from MINFLUX data (*MT-ND1*, *MT-CO3*, and *MT-CYB* mRNAs). The analysis is based on the same data as in Fig. 2F. **(B)** 3D rendition of combined MINFLUX localizations of all 13 mRNAs (yellow) and the ribosomal protein bL12m (cyan). **(C)** Confocal images of U-2 OS cells: control cells treated with scrambled siRNA and U-2 OS bL12m knockdown cells (KD) cells treated with siRNA pools for eight days. Cells were immunolabeled with the same anti-bL12m antibody as used for MINFLUX (upper panels) and an anti-ATPB antibody (lower panels). The imaging settings and contrast settings were identical for control and knockdown cells, demonstrating the specificity of the anti-bL12m antibody. **(D)** Close-ups of several sites from (B). Each sphere represents the combined MINFLUX localizations of a single fluorophore and was rendered to have a diameter of 10 nm. Box: 25/75 percentile; whiskers: max/min without outliers; line: median; square: mean. Curved line: kernel density estimation representing the data point distribution (A). Scale bars: 1 µm (C). Sphere diameter = 10 nm (B, D).

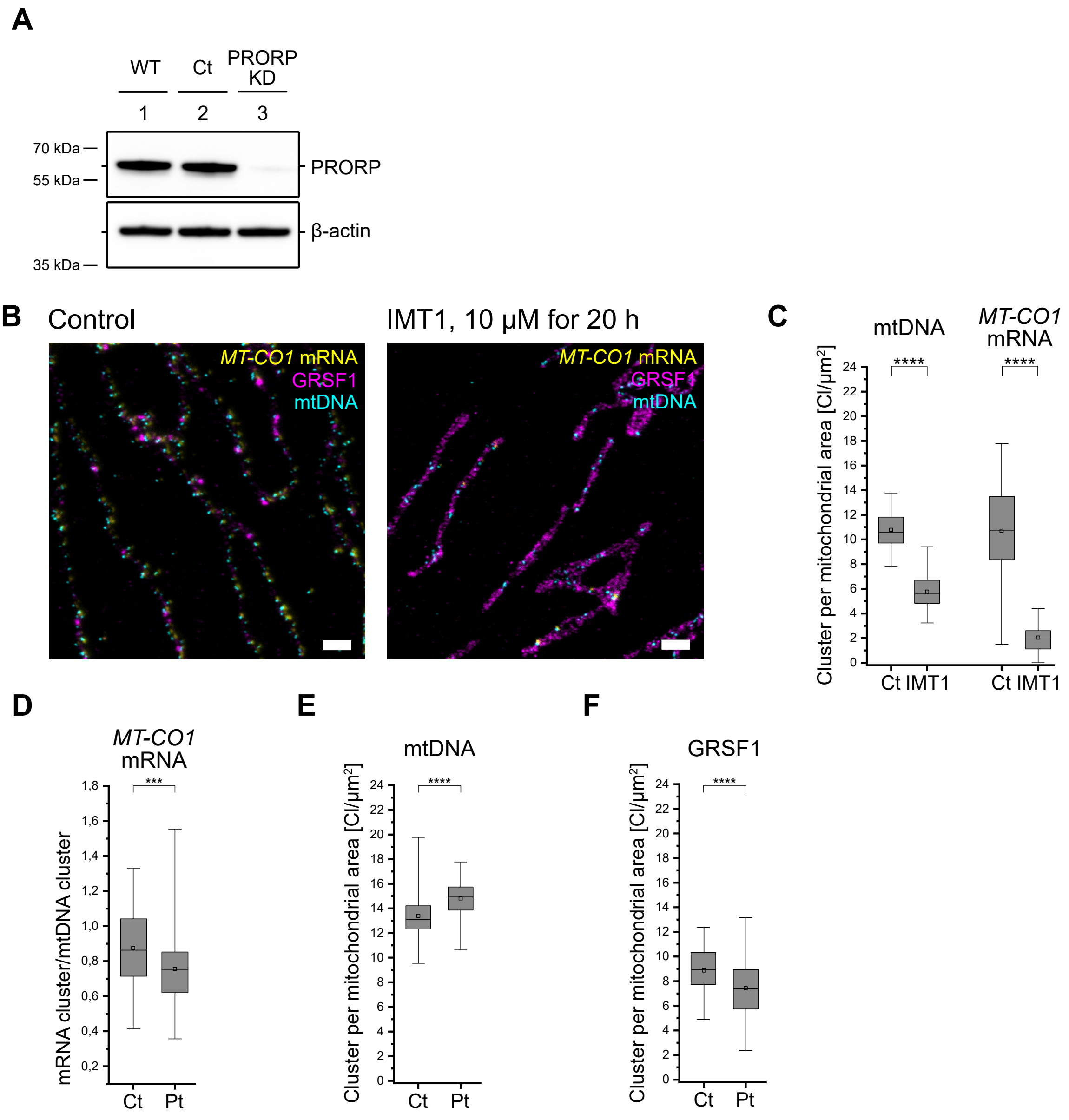


### Suppl Figure 4: STED-smFISH enables the investigation of mRNA distribution changes

**(A)** Western blot of whole cell lysate of untreated U-2 OS cells (WT), U-2 OS cells treated with scrambled siRNA (Ct) and U-2 OS PRORP knockdown (PRORP KD) cells treated with siRNA pools for six days, decorated with an anti-PRORP antibody and an anti-beta-actin antibody. **(B)** STED images of U-2 OS cells labeled for GRSF1 (magenta), mtDNA (cyan), and *MT-CO1* mRNA (yellow). The cells were either untreated (control, left) or treated with 10 µM IMT1 for 20 h to inhibit mitochondrial transcription (right). **(C)** Quantification of mtDNA and *MT-CO1* mRNA clusters per mitochondrial area in untreated control (Ct) and IMT1 treated cells (n=66 images (Ct), n=71 images (IMT1)). **(D)** Quantification of the *MT-CO1*-mRNA cluster to mtDNA cluster ratio in control fibroblasts (Ct) and fibroblasts from a patient (Pt) carrying the m.14709T>C mutation in gene MT-TE (for image data see Fig. 3E). **(E)** Quantification of mtDNA clusters per mitochondrial area in control fibroblasts (Ct) and fibroblasts from a patient (Pt) carrying the m.14709T>C mutation in gene MT-TE (for image data see Fig. 3E). **(F)** Quantification of GRSF1 spots per mitochondrial area in control fibroblasts (Ct) and fibroblasts from a patient (Pt) carrying the m.14709T>C mutation in gene MT-TE (for image data see Fig. 3E). Box: 25/75 percentile; whiskers: max/min without outliers; line: median; square: mean; ****: p-value ≤0.0001; ***: p-value ≤0.001 (C-F). Scale bars: 1 µm (B).

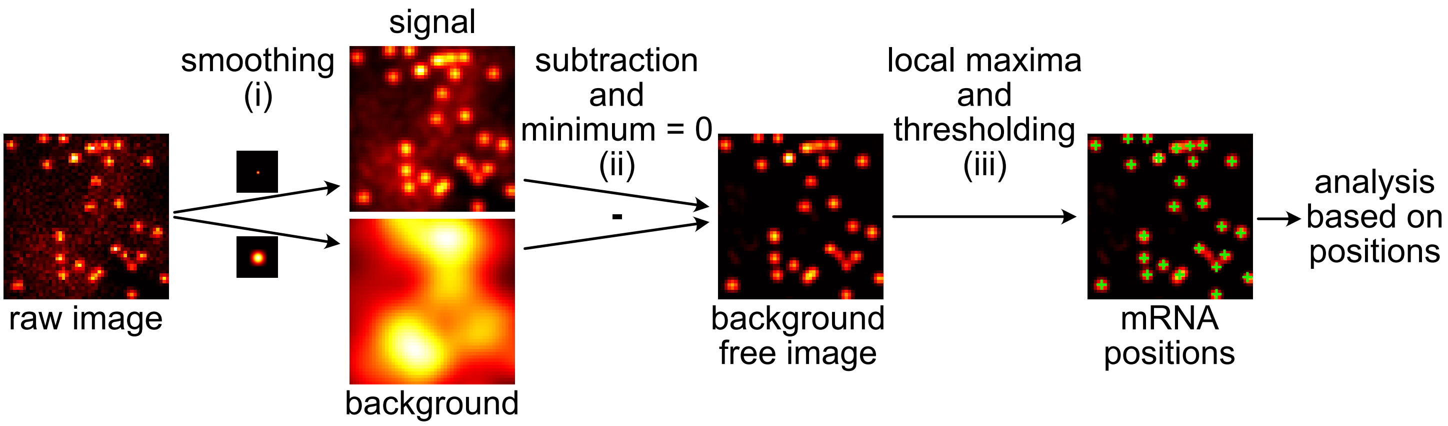


Suppl Figure 5: STED-smFISH image data analysis

Analysis pipeline for determining mRNA positions from STED-smFISH image data. The raw images are smoothed with a 25 nm FWHM Gaussian filter (signal) and a 210 nm FWHM Gaussian filter (background) (i). The background is subtracted from the signal and the minimum in the images is set to zero (ii). Finally, the local maxima with at least 3 counts were detected (iii) and used as mRNA positions. The positions can be used for subsequent analysis.
